# Smart Lattice Light Sheet Microscopy for imaging rare and complex cellular events

**DOI:** 10.1101/2023.03.07.531517

**Authors:** Yu Shi, Jimmy S. Tabet, Daniel E. Milkie, Timothy A. Daugird, Chelsea Q. Yang, Andrea Giovannucci, Wesley R. Legant

## Abstract

Light sheet microscopes enable rapid, high-resolution imaging of biological specimens; however, biological processes span a variety of spatiotemporal scales. Moreover, long-term phenotypes are often instigated by rare or fleeting biological events that are difficult to capture with a single imaging modality and constant imaging parameters. To overcome this limitation, we present smartLLSM, a microscope that incorporates AI-based instrument control to autonomously switch between epifluorescent inverted imaging and lattice light sheet microscopy. We apply this technology to two major scenarios. First, we demonstrate that the instrument provides population-level statistics of cell cycle states across thousands of cells on a coverslip. Second, we show that by using real-time image feedback to switch between imaging modes, the instrument autonomously captures multicolor 3D datasets or 4D time-lapse movies of dividing cells at rates that dramatically exceed human capabilities. Quantitative image analysis on high-content + high-throughput datasets reveal kinetochore and chromosome dynamics in dividing cells and determine the effects of drug perturbation on cells in specific mitotic stages. This new methodology enables efficient detection of rare events within a heterogeneous cell population and records these processes with high spatiotemporal 4D imaging over statistically significant replicates.

## Introduction

Light sheet microscopes offer reduced photobleaching, less phototoxicity, and increased imaging speed compared to widefield or confocal microscopes^1,2^. These advantages span spatiotemporal scales, permitting the visualization of dynamic events ranging from single molecules that diffuse in milliseconds ^3–7^ to organisms that develop over multiple days^8–11^. However, these advances also present new challenges. First, light sheet microscopes can extract gigabytes per second of information from the specimen which necessitates specialized data storage, visualization, and quantification tools^12^. Second, biological samples can rarely tolerate hardware-limited maximum imaging rates for very long before becoming perturbed by the imaging process^13^. Third, because data is still acquired plane-by-plane, there is an inherent sacrifice between the volumetric sampling rate and the imaging field of view driving a tradeoff between rapid, high-content imaging of a small region vs high-throughput imaging of many samples or replicates. Lastly, the oblique geometry, high-magnification, and limited field of view used by many light sheet microscopes impede the rapid selection of specific cells or regions of interest across a large sample, especially if such cells are rare within a population^6,14–16^.

By detecting and responding to specific image features, event-triggered microscopy has emerged as a way to overcome these tradeoffs. Previous demonstrations include locating rare cells within a population and automating 3D confocal imaging experiments^17^, selectively increasing the acquisition rate to capture mitochondrial fission and bacterial cell division events with structured illumination microscopy^18^, and switching between diffraction limited and super-resolution stimulated emission depletion modalities when specific spatial signatures are detected within live-cell time-series^19^. Inspired by these recent advances, we sought to determine whether similar approaches could be used to program a self-driving lattice light sheet microscope “smartLLSM” that is capable of searching for specific cells within a population and then automatically capturing 4D light sheet movies of these specimens. We accomplish this by rapidly scanning centimeter-scale specimens via inverted epifluorescent microscopy and switching on demand, to perform lattice light sheet microscopy (LLSM) volume imaging at automatically determined fields of view. We automate the instrument by feeding images to a fully convolutional machine learning algorithm “You Only Look Once, version 5 (YOLOv5)” for real-time object detection and classification^20,21^. The locations and classes of identified objects feedback to the instrument to automate imaging of desired cells within a population. We chose to use fluorescently labeled nuclei and cell division as a benchmark application because the cell stage can be visually determined by chromosome morphology, and because it is a highly dynamic 3D process that substantially benefits from the low-phototoxicity imaging afforded by light sheet microscopy^22–24^. This approach processes hundreds of cells/second dramatically cutting down user interaction time and user bias in sample selection to generate statistically significant, high-content replicates of rare cellular events. We combine this method with 3D/4D image analysis to quantify mitotic spindle orientation, kinetochore position, and chromosome morphology at different mitotic stages and to longitudinally track kinetochore trajectories throughout mitosis. The automated nature of this approach allows a user to capture hundreds of 3D images or dozens of 4D movies and to quantify subtle drug perturbation effects, even for rare and transient cellular stages. Overall, smartLLSM optimally balances the competing needs of high-content light sheet microscopy with the need to capture statistically significant replicates of rare cellular events. This allows it to dramatically reduce the need for up-front data storage and post-acquisition data mining to draw meaningful biological conclusions.

## Results

### smartLLSM incorporates a fast and accurate object detector

To create training data, we constructed a retinal pigment epithelium cell line stably expressing fluorescent fusions of centromere protein A (CENPA-mNeonGreen) and histone H2B (H2B-mScarlet) to label centromeres/kinetochores and chromosomes respectively. Using this cell line, we generated annotated training data via inverted epifluorescent microscopy of the chromosome channel (**Fig. 1A i**). Nuclei were segmented with the deep-learning based algorithm Cellpose^25^ and regions of interest around each cell were cropped and annotated according to cell stage (**Fig. 1A ii**). Because over 90% of the cells are in interphase, to facilitate manual annotation of the more rare cell stages, we utilized a bootstrap approach to first train a three-state classifier for cells in interphase, mitosis, and a “blurry” class to deal with out of focus regions or non-specific cellular debris (**Fig. 1A iii, Fig. S1**). Subsequently, the subset identified as mitotic cells was further split into 6 cell classes (prophase, prometaphase, metaphase, anaphase, and telophase) (**Fig. 1A iv**). In total, we annotated ~54,000 cells with approximately 5% comprising different mitotic stages. Finally, the raw images, the bounding boxes determined via Cellpose, and the class for each cell were used to train YOLO (**Fig. 1A v**). Our trained model achieved an average accuracy of 85% for all mitotic stages (**Fig. 1B**). Precision vs. recall curves, calculated as “one vs. all” for each class, can be used to tune the model to maximize the network’s precision (low false positive rate) or recall (low false negative rate) (**Fig. 1C**).

**Figure 1:**
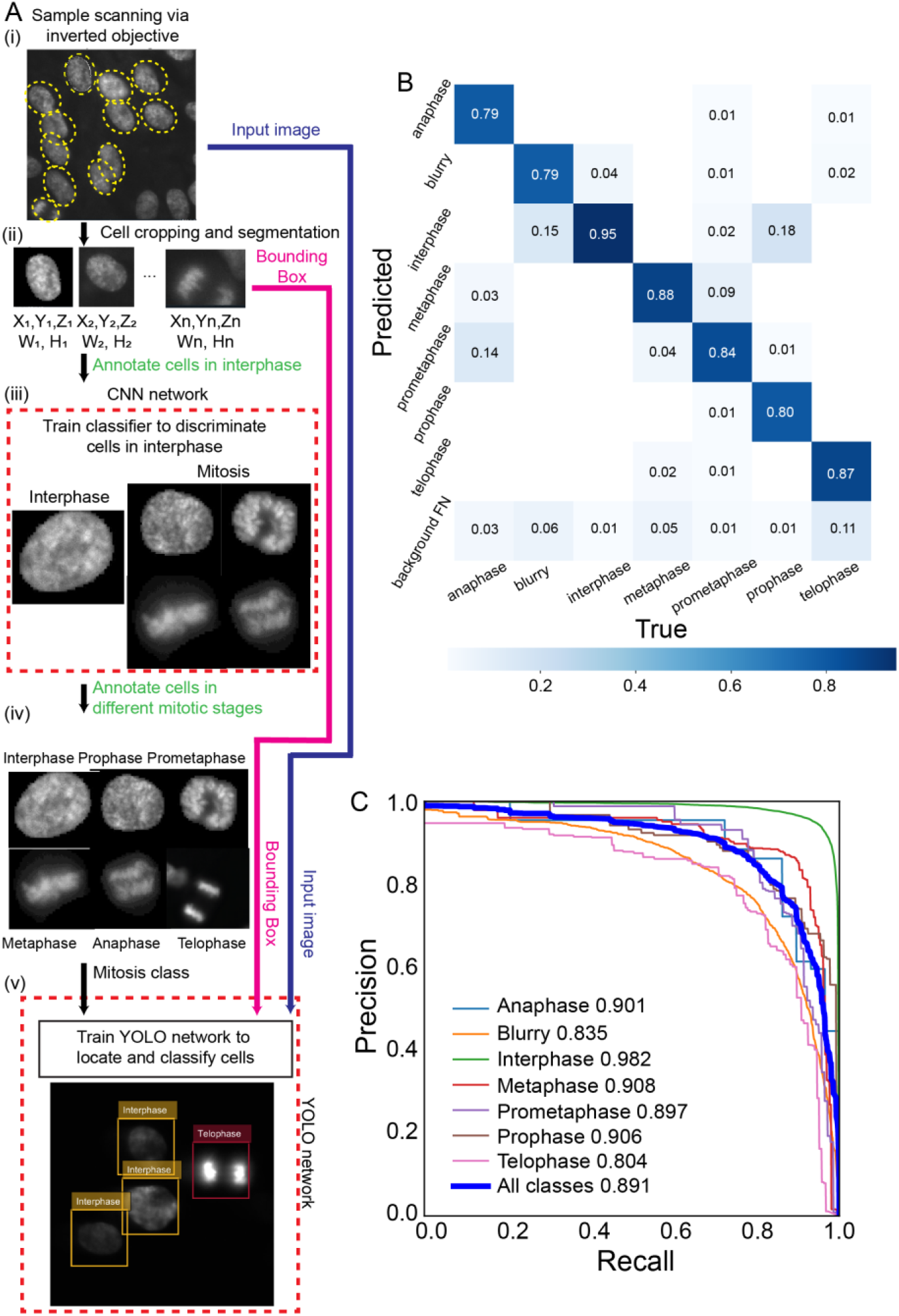
Training and validation for smartLLSM. **(A)** Schematic of the smartLLSM training process. Cell nucleus images taken with the inverted objective (i) were segmented by Cellpose to extract each cell’s location and nucleus mask size (ii). A portion of the segmented cells were manually annotated to distinguish cells in interphase from mitosis and the results were used to train a convolutional neural network to filter images with a high likelihood of containing mitotic cells (iii). Cells within each image were manually annotated by their stage (iv), and the raw image (blue), the location and size of the bounding box (magenta), and the associated class (including interphase and different mitotic stages) were then fed into training a YOLOv5 network (v). **(B)** Confusion matrix for test images across all detected classes. (C) Precision vs. recall curves for each detected class. The numbers in the legend indicate the area under each curve.

### smartLLSM implementation and timing

To demonstrate the utility of smartLLSM, we highlight use cases for both fixed and live-cell imaging. In each of these cases, we perform a 2D tiled scan of a 16 mm x 6 mm area using inverted epifluorescent microscopy while passing the tiled images to the YOLOv5 network (**Fig. 2A**). For a typical dataset, we process batches of 100 images in parallel, with each image consisting of 800 x 800 pixels. With a pixel size of 100 nm at the sample, each individual image is an 80 x 80 μm field of view and each batch represents a 10 x 10 tiled array of images from 800 x 800 μm of the sample. When plated at 80% cell confluence, each image batch contains approximately 300 nuclei and requires 1.4 seconds to process (**Fig. S2**). This yields a network processing rate of ~214 cells per second dramatically exceeding the rate of manual image inspection.

**Figure 2:**
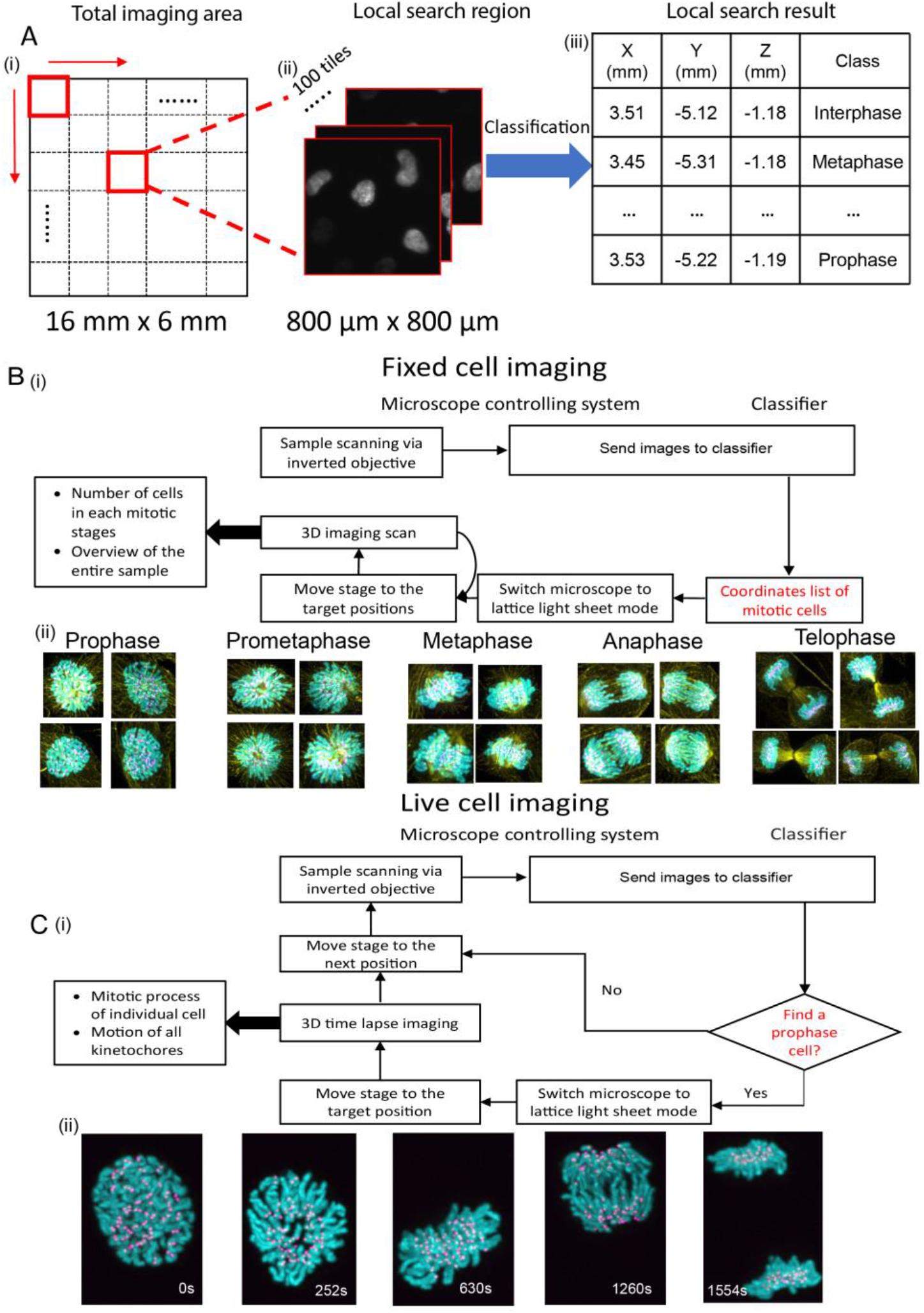
smartLLSM workflow. **(A)** Flowchart for smartLLSM imaging. A total imaging area of 16 mm by 6 mm is segmented into a grids of 800 μm by 800 μm squares (i). At each location, 100 images of 80 μm by 80 μm fields of view are imaged via inverted epifluorescence microscopy (ii). These images are sent to the YOLOv5 network to extract the location and associated mitotic class for each cell (iii). **(B)** Flow chart for imaging of fixed samples. After scanning is complete, the microscope switches to LLSM imaging to acquire a gallery of 3D image scans for each cell stage of interest (i). Images show chromosomes (cyan), kinetochores (magenta), and microtubules (yellow) (ii). **(C)** Flow chart for imaging live-sample dynamics. Once a cell in prophase is found, the microscope switches to LLSM imaging to capture a 4D movie of the mitotic process (i). Images show representative time points from a live-cell movie of kinetochores (magenta) and chromosomes (cyan) (ii).

For fixed specimens, we scan the entire area via 2D inverted epifluorescent microscopy while logging the stage and position of each cell (**Fig. 2B i**). With a camera exposure time of 50 ms, the entire 96 mm^2^ imaging region containing approximately 30,000 cells can be scanned with 300 nm resolution in ~15 minutes while processing the resulting 11,400 2D images takes ~10 minutes for cell detection and classification including file input/output overhead. Pipelining image acquisition and network inference in parallel allows for real time processing. Once complete, we then switch to LLSM and return to each cell of interest to perform multicolor 3D imaging (**Fig. 2B ii**). Scanning a 60 μm by 110 μm by 30 μm volume around each cell at 50 ms exposure per plane with three colors via LLSM takes approximately 20 s, although this could be sped up by an order of magnitude or more by shortening the exposure-per-plane, depending on fluorophore brightness and desired imaging quality. Altogether, this use case provides automatic and high-throughput 3D imaging of selective cells within a population (**Movie S1**).

In the second use case, we continuously scan living cells via 2D inverted epifluorescent microscopy until we identify a cell in prophase. Once a prophase cell is found, a feedback loop drives the stage to the target coordinates (**Fig. 2C i**) and the microscope is switched to lattice light sheet mode. If multiple prophase cells are detected within a single image, then the cell with the highest score is given priority. After centering the cell, the microscope collects a 50-minute lattice light sheet volume time-series, which for unperturbed cells is sufficient to capture the entire mitotic process (**Fig. 2C ii, Movie S2)**. After collecting the time-series, the microscope then switches back to 2D inverted image scanning until the next prophase cell is detected (**Fig. 2C i**). The timing for the inverted imaging and classification are the same as for the fixed cell case above. This demonstrates the capability for automated, event-triggered 4D imaging of rare and transient biological phenomena like cell division.

### smartLLSM to surveil cell populations

To verify that that the trained YOLO network performed as expected, we sought an orthogonal validation that did not rely on user interpretation of mitotic state. To this end, we determined the proportion of mitotic cells found by our network when imaging a synchronized cell population compared to a control sample. Under live-cell conditions, we repeatedly imaged a 16 mm x 6 mm area via inverted epifluorescent microscopy every 30 minutes while feeding the images to our classification network. Fluorescent RPE reporter cells, synchronized by a double thymidine block, showed a clear peak in the percent of mitotic cells between 10 and 14 hrs after thymidine release whereas this peak was absent in unsynchronized control samples (**Fig. 3A**). Furthermore, in control samples the ratio of mitotic cells increased slightly over the course of the 18 hr imaging experiment, indicating minimal phototoxicity or perturbation from being maintained in the microscope imaging chamber or from the imaging itself.

**Figure 3:**
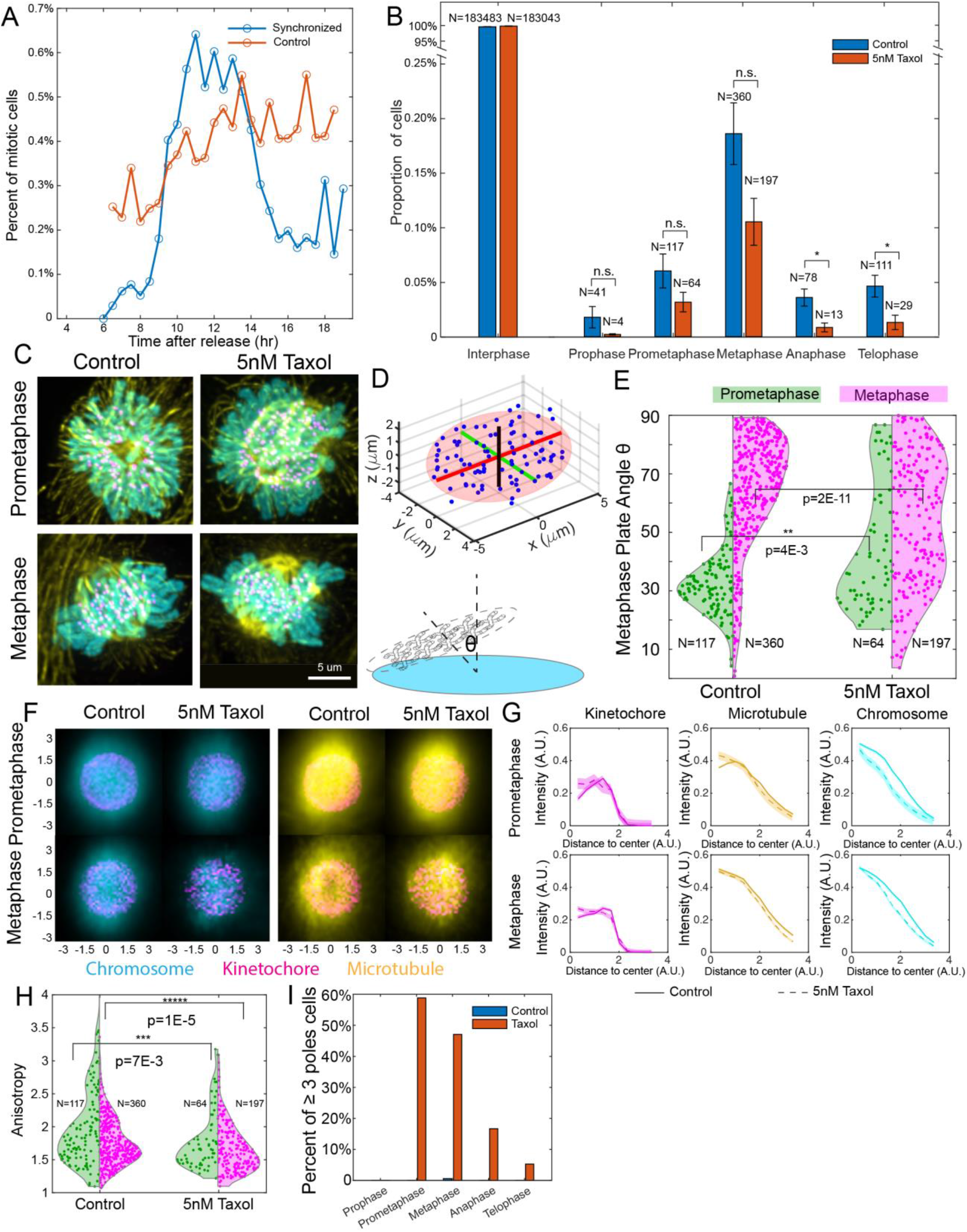
High throughput imaging of Taxol induced mitotic defects. **(A)** A plot of the percentage of mitotic cells in control (orange) and synchronized (blue) samples over time after thymidine block release showing mitotic wave detection. **(B)** Proportion of cells in different mitotic stages for control (blue) and 5 nM Taxol treated (orange) RPE cells. The number above each bar indicates the total number of cells in the corresponding stage. The data is integrated over three independent samples for each condition, the bar plot and the error bars indicate the mean and the standard error of the mean respectively. *:p<0.05 **(C)** Representative maximum intensity projections (MIPs) of prometaphase and metaphase under control and 5 nM Taxol treated conditions showing chromosomes (cyan), kinetochores (magenta), and microtubules (yellow). **(D)** Diagram for the characterization of kinetochore distributions. Top: anisotropy is defined by the ratio between the variation of the axis with the largest and the smallest variation (see Methods for details). Bottom: Diagram depicting the metaphase plate orientation relative to z-axis of the microscope objective (see Methods for details). **(E)** Violin plot of metaphase plate angle (θ) for control and 5 nM Taxol treated cells in prometaphase (green) and metaphase (magenta). The colored areas indicate the distribution. Horizontal bars indicate the significance based on the Kolmogorov–Smirnov test (K-S test) between Taxol and control in prometaphase (N = 117 for control and N = 64 for Taxol) and metaphase (N = 360 for control and N = 197 for Taxol). **(F)** Averaged heatmaps of kinetochore (magenta), chromosome (cyan), and microtubule (yellow) distributions for prometaphase and metaphase cells under control and 5 nM Taxol treated conditions. **(G)** Radial intensity profiles for the labels shown in (F). The line and shades indicate the mean and the standard error of the mean, respectively at each distance. Solid lines indicate control, and dashed lines indicate 5 nM Taxol treated condition. **(H)** Violin plot of anisotropy for control and 5 nM Taxol treated cells in prometaphase (green) and metaphase (magenta). The horizontal bars indicate the K-S significance test between the control and 5 nM Taxol-treated condition in prometaphase and metaphase cells. **(I)** The ratio of >=3 pole spindles in different mitotic stages in control (blue) and 5 nM Taxol-treated (orange) conditions

We next sought, to demonstrate the advantages of smartLLSM for balanced high-throughput and high-content imaging of select cells. To do so, we imaged chemically fixed RPE cells expressing CENPA-mNeonGreen and H2B-mScarlet. Across multiple coverslips, we located and classified approximately 180,000 cells of which only, ~1 percent were in mitosis, which is similar to our live-cell control samples above (**Fig. 3B**). Of the mitotic cells, the majority were in metaphase (0.195%) while prophase was the rarest sub-class (0.022%). This demonstrates the challenge of studying cells in prophase at high resolution while also obtaining sufficient statistical replicates to test the effect of a perturbation. A user would need to manually scan through roughly 4,500 cells before finding a single cell in prophase to image. Using smartLLSM dramatically accelerates this process.

### smartLLSM for high-content and high-throughput imaging of specific cell stages

SmartLLSM allows a user to automatically capture both population-level statistics on cell stage and high-resolution images of select cells within that population. To demonstrate the utility of this approach, we compared samples with and without treatment with Paclitaxel (Taxol), a commonly used chemotherapeutic. Previous work has indicated that low-concentration, clinically-relevant Taxol treatment of 5 nM delays cell mitosis^26^ and disrupts cell survival^27^ by stabilizing microtubules. Both control and Taxol treated conditions had the vast majority of cells in interphase, but Taxol treatment significantly reduced the percentage of cells in different mitotic stages (p = 0.02) (**Fig. 3B**). We also verified that Taxol treatment altered the distribution of mitotic cells across different stages using a chi-square test of independence (p = 7E-8) with significant reductions in the portion of anaphase and telophase cells (p < 0.05). This implied that, as expected, Taxol-treated cells are inhibited progressing through downstream mitotic stages. To quantify the sub-cellular effects of Taxol perturbation on specific mitotic stages, we automatically imaged kinetochores, H2B, and microtubules (immunostained for β-tubulin-Alexa647) via smartLLSM on the same coverslips, focusing in more detail on cells in prometaphase and metaphase (**Fig. 3C**). We first transformed 3D LLSM datasets of different cells to the same reference frame using the kinetochore distribution and the plane of the metaphase plate as a coordinate origin (see Methods for details). When we examined the orientation of the metaphase plate relative to the cover glass, we found a clear reorientation from parallel (in prometaphase) to perpendicular (in metaphase) (**Fig. 3D, E**), in agreement with previous observations tracking the spindle pole orientation during these stages^28^. Interestingly, this reorientation was disrupted by exposure to Taxol, leading to a randomly orientated metaphase plate (**Fig. 3E**). Previous studies also reported that shortly after nuclear envelop breakdown, kinetochores and chromosomes form a ring-like structure in a zone that is enriched for spindle microtubules^28,29^. It is thought that this structure facilitates accurate chromosome segregation, increasing the probability of correct amphitelic attachments between kinetochores and microtubules by providing increased lateral attachment contacts between kinetochores to spindle-associated microtubules^28^. Therefore, we sought to determine whether we could replicate this result and also test the effects of low-dose taxol treatment on this early-stage mitotic structure. By averaging the kinetochore, chromosome and tubulin signals across cells, we found a clear ring-like kinetochore distribution that was maximal in prometaphase and was persistent into metaphase (**Fig. 3 F, G**). At prometaphase, the maximal microtubule density coincided with the peak of the kinetochore distribution, replicating the “barrel-like” structure observed in prior studies^30^, whereas at metaphase, the peak microtubule intensity was at the center of the metaphase plate. However, in contrast to previous studies, we did not find that, on average, chromosome bodies were depleted within the ring core in prometaphase classified cells^28,29^. This may indicate that this is a transient stage in prometaphase or that not all cells display such a structure. Interestingly, Taxol treatment yielded a less-prominent kinetochore ring structure at both prometaphase and metaphase and increased the microtubule density at the core of the kinetochore ring (**Fig. 3F, G**).

To better understand these results, we quantified the anisotropy of kinetochore distributions in prometaphase and metaphase cells. As expected, spatial anisotropy increased as the kinetochores assembled into an ordered metaphase plate and was decreased at both prometaphase and metaphase by treatment with Taxol (**Fig. 3H**). Examining the microtubule distribution from the 3D LLSM images revealed that this was likely due to an increase in the presence of multipolar (≥3) spindles in Taxol treated cells. These multipolar structures were not present in prophase cells, but peaked to nearly 60% of all cells examined in prometaphase before declining gradually for other mitotic cell stages (**Fig. 3I**). In summary, targeted acquisition of selected cell populations via smartLLSM overcomes the competing demands of high-content and high-throughput imaging of cells in mitosis. Our studies reveal the subtle effects of low-dose pharmacological perturbation on specific stages of the mitotic process and illustrate potential of this approach for future imaging applications in drug screening on specific cells or stages within a population.

### smartLLSM for automated 4D imaging of cell division

Fixed cell imaging provides only a snapshot of a dynamic 3D process. As such, it is difficult to capture transient temporal events and it is impossible to quantify longitudinal changes in dynamic properties within cells. To address this need, we utilized smartLLSM for event triggered live-cell imaging of cell division utilizing the dual-labeled reporter line described above starting with prophase and continuing through cytokinesis. Mitosis is an inherently dynamic event that requires rapid 3D imaging. However, mitotic cells are also especially sensitive to perturbation from phototoxicity, often arresting in prometaphase if light doses are too intense^13^. As a first application, we utilized smartLLSM to capture a number of movies under different imaging conditions and to optimize the light dose, exposure time, and imaging rate while ensuring proper cell division (**Table S1**). In early prophase, kinetochore motion is highly intermittent and requires rapid imaging to unambiguously track individual kinetochores. However, fast imaging also leads to increased photobleaching which degrades the ability to detect kinetochores at later mitotic stages. To optimally capture these dynamics while preventing long-term photobleaching, we imaged in an adaptive manner, first imaging at a rate of 3 seconds/volume for the first 5 minutes and then transitioning to a rate of 6 seconds/volume for the following 45 minutes of imaging. From these movies, we studied the motion of kinetochores by longitudinally tracking them in three dimensions ^31^ (**Movie S3**).

We first examined the instantaneous speed of kinetochores from prophase to metaphase by comparing their locations across time points. During the transition from prophase to prometaphase, we noticed that kinetochores show a rapid intermittent inward movement which occurs shortly after nuclear envelop breakdown and during the initial stages of kinetochore capture by microtubules^24^. Based upon this, we partitioned each live-cell volumetric movie and plotted the kinetochore speed distribution for each of four stages: prophase, contracting, prometaphase, and metaphase, (**Fig. 4A, Movie S3**). This data revealed an increase in kinetochore speed after nuclear envelop breakdown in prophase, but no significant differences between the contracting, prometaphase or metaphase stages. In all cases, kinetochore speed was well above our noise floor of 10 nm/s, estimated by repeatedly imaging chemically fixed cells at different stages (**Fig. S3**). We next investigated whether kinetochore speed might be differentially sensitive to microtubule dynamics during specific mitotic stages by examining the effects of Taxol treatment on these measurements (**Fig. 4B, Movie S4**). To test this, we first screened a range of Taxol concentrations for their effect of the cells ability to complete mitosis within a 50-minute time window. Cells treated with 5 nM Taxol were still able to enter prophase. However, compared to control cells, none of the cells imaged under 5 nM Taxol conditions progressed through mitosis (33/35 vs. 0/27), typically arresting at the prometaphase to metaphase transition. In contrast, cells treated with 0.5 nM Taxol progressed similarly to control (16/17). Cells treated with 1 nM Taxol lay in between, with roughly one third of the cells completing mitosis and two thirds arresting prior to metaphase (7/23) (**Fig. S4**). When we observed the effects of Taxol dose on kinetochore velocities (**Fig. 4B**), we found that 5 nM Taxol strongly reduced kinetochore speed in prometaphase compared to control cells. For 1 nM Taxol treated cells that completed mitosis successfully, there was no significant difference in kinetochore speed at any cell stage, whereas for those that eventually arrested, we observed decreased kinetochore speed during prometaphase with an intermediate magnitude of effect compared to the 5 nM dose. Taxol treatment had no effect on kinetochore speed during prophase which was expected given this stage is prior to nuclear envelop breakdown. However, kinetochore speed during the initial contracting stage of kinetochore motion after nuclear envelop breakdown was less sensitive to Taxol than kinetochore speed during prometaphase (**Fig. 4C**), suggesting that other factors besides microtubule dynamics may impact kinetochore motion during this stage. Live kinetochore tracking also revealed that the metaphase plate reorientation between prometaphase and metaphase images of fixed cells occurs within the first 400 seconds of cell division after prophase, but is delayed and diminished under Taxol treated conditions (**Fig. 4D, Fig. S5**).

**Figure 4:**
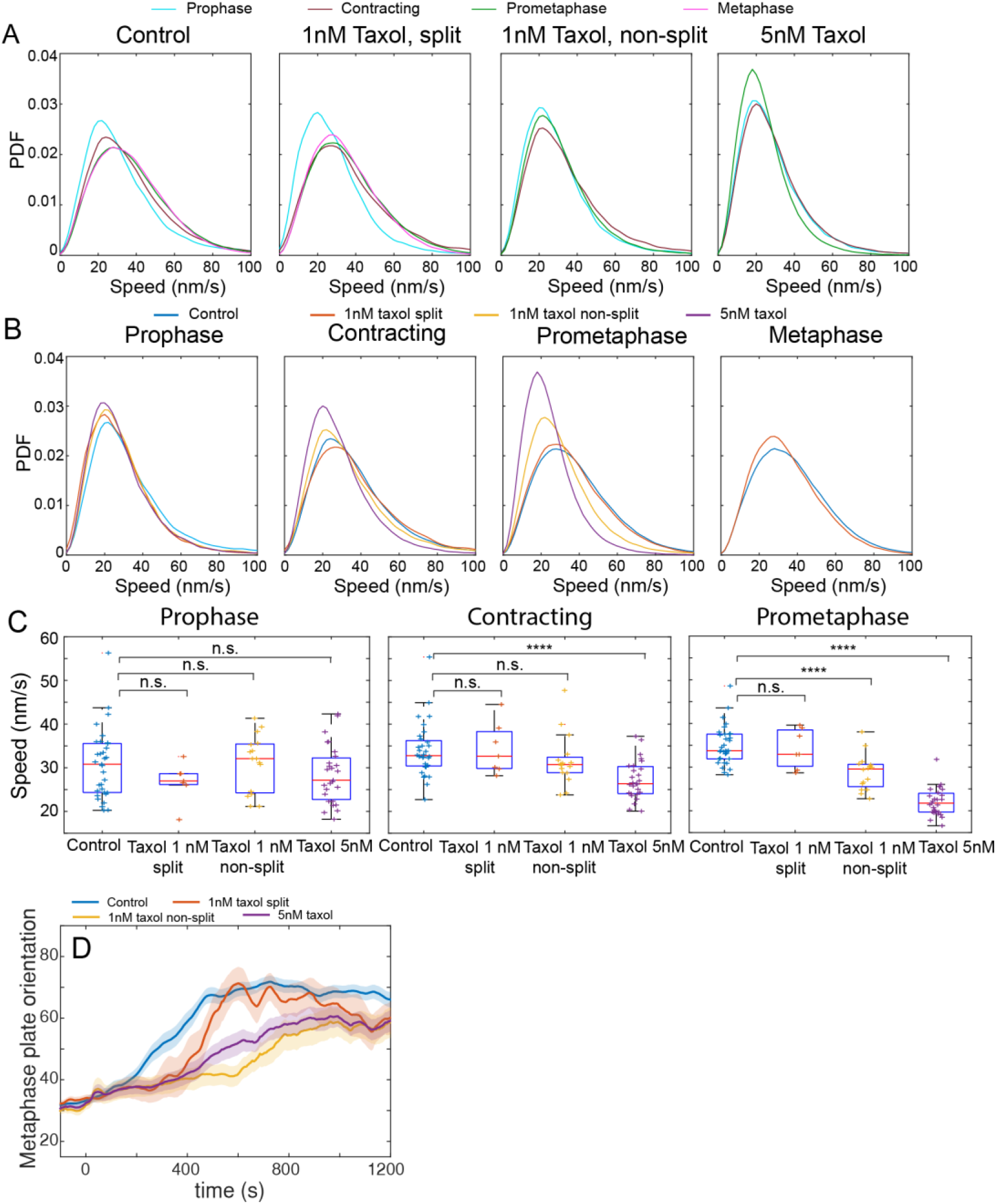
High throughput imaging of kinetochore dynamics during mitosis. **(A)** Distribution of kinetochore speed during various mitotic phases under different conditions. Four different mitotic stages are included: prophase (cyan), contracting phase (dark red), prometaphase (green), and metaphase (magenta). Four different conditions are investigated: control (N = 35), 1 nM Taxol treated cells that finish mitosis (N = 7), 1 nM Taxol treated cells that fail to divide (N=16), and 5 nM Taxol treated cells (purple, N = 27). **(B)** Distribution of kinetochore speed similar to (A) but grouped by mitotic stage. **(C)** Box and whisker plot for the median kinetochore speed of a given cell at a given stage. The box indicates the 25^th^ to 75^th^ percentiles and the horizontal marker indicates the population median. The whiskers extend to the most extreme data that are not considered outliers. All perturbed conditions are compared to the control condition via t-test. n.s.: not significant, ****:p<1E-4. **(D)** Metaphase plate orientation (θ) plotted over time for each cell. The solid line indicates the average angle at a given time after the start of prometaphase (t = 0 s, see Methods for details) and the shades indicate the standard error across cells.

### smartLLSM for longitudinal kinetochore tracking in mitosis

In addition to quantifying instantaneous kinetochore speed, smartLLSM also allows for longitudinal tracking of kinetochores throughout the mitotic process. Prior studies have recently used LLSM and 3D single particle tracking to identify “lazy” or lagging kinetochores which fail to partition into the daughter cells at the same rate during anaphase^32^. Intriguingly, this study revealed that metaphase kinetochore dynamics could be partially predictive of future mislocalizations during anaphase. However, measurements were limited by the ability to only track kinetochores from metaphase to anaphase. Here, we demonstrate the utility of smartLLSM to automatically image and track kinetochores throughout mitosis and identify outlier kinetochores based on their spatial coordinates relative to the overall distribution of kinetochores within the cell at a given time point (see Methods for details on outlier identification). Once an outlier kinetochore is identified, its trajectory can be analyzed to reveal its behavior before, during, and after it deviated from the population distribution (**Fig. 5A, Movie S5**). Although we did not observe any significant change in the speed of outlier kinetochores vs. others (**Fig. S6**), we found that the probability of observing an outlier kinetochore varied at different mitotic stages (**Fig. 5B**). Since chromosomes are contained within the nuclear envelop during prophase, we did not detect any outliers at this stage, but detected that approximately 1% of kinetochores, measured across 33 cells, were outliers in both prometaphase and anaphase. In contrast, only 0.2% of kinetochores are outliers in metaphase (**Fig. 5B**). Chromosome positions undergo the largest reorganizations during prometaphase and anaphase, so the potential for mitotic errors and for kinetochores to deviate from the population average may be higher in these stages than during metaphase when most chromosomes move relatively little within the metaphase plate. We next examined average duration when a kinetochore is classified as an outlier to the time at which it no longer was considered an outlier. Interestingly, outlier events were longest in metaphase (238.6 s), followed by prometaphase (505.8 s) and shortest in anaphase (78.4 s), indicating that the mechanisms for outlier correction may take longer during metaphase (**Fig. 5C**). By considering both the number of outlier kinetochores in a given stage, the duration of the outlier events themselves, and the average duration of a mitotic stage, we can compute the instantaneous probability of observing an outlier kinetochore at any instant in time within given cell stage. With this information, we find that at any given time, there is an approximately 0.26%, 0.14%, and 0.20% percent chance to observe an outlier kinetochore in prometaphase, metaphase, and anaphase respectively (**Fig. 5D**).

**Figure 5:**
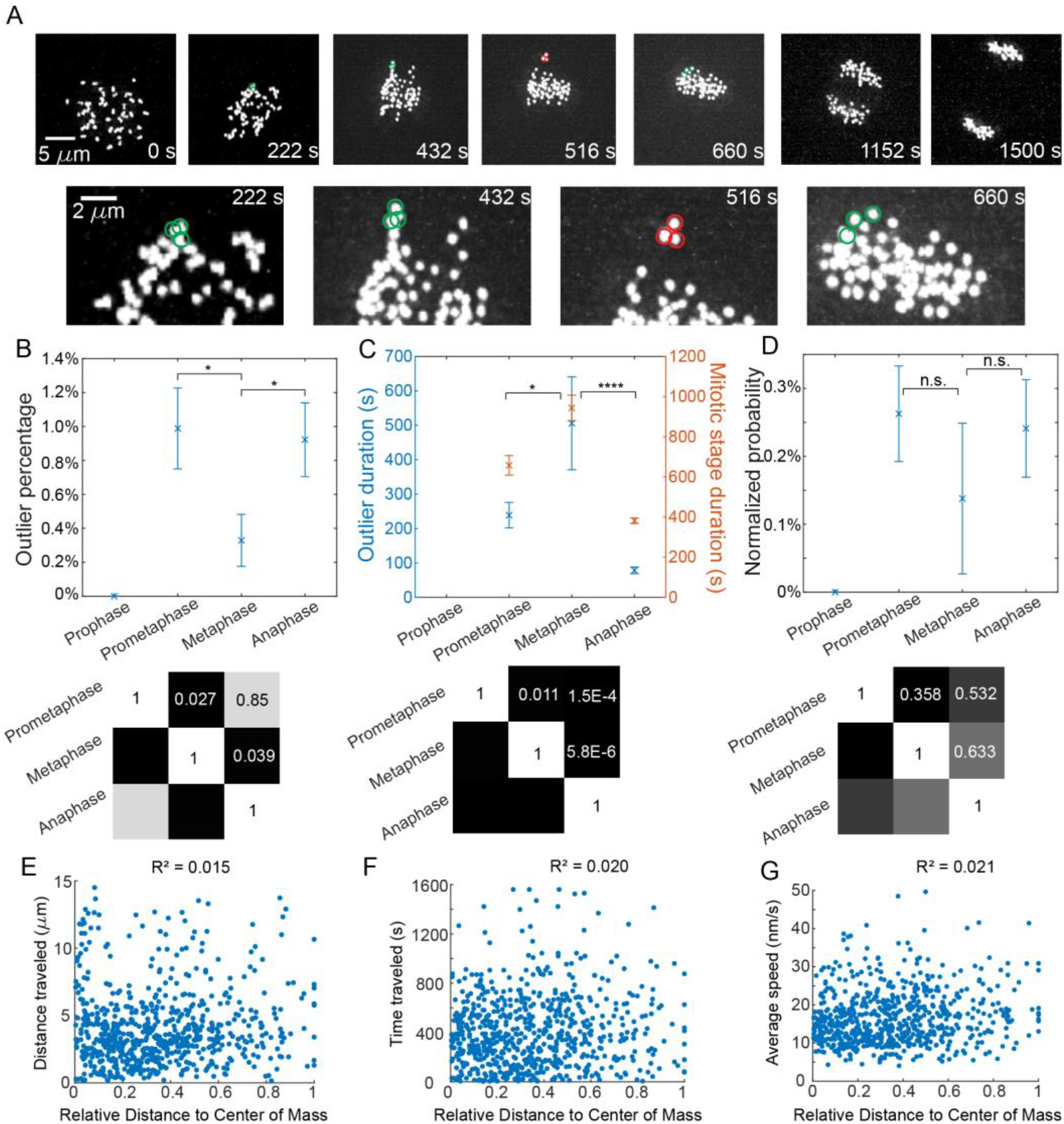
Longitudinal tracking of kinetochore motion during mitosis. **(A)** Sample snapshots of the kinetochore trajectories during mitosis. T = 0 s is the start of the movie in prophase. Red circles indicate kinetochores that are identified as outliers. Green circles indicate the locations of outlier kinetochores prior to and after being identified as outliers. **(B)** Plot of the percentage of kinetochores that are considered an outlier in different mitotic stages measured across N = 33 control cell movies. Crosses and error bars indicate the median and the standard error measured across individual cell movies. **(C)** Plot of the durations of outlier events (blue, left axis) and the durations of different mitotic stages (orange, right axis). The crosses and error bars indicate the mean and the standard error of outlier kinetochores detected in different stages: prophase N = 0, prometaphase N = 30, metaphase N= 10, anaphase N = 28. Here N is the number of outlier kinetochores across all 33 control cell movies. **(D)** Plot of the instantaneous probabilities for finding an outlier kinetochore at a given time point in different mitotic stages (see Methods for details). The matrices below B-D show the p-value of the pairwise K-S significant test between prometaphase, metaphase, and anaphase. **(E-G)** Scatter plots of kinetochore location within the prophase nucleus (distance to the center of mass of the kinetochore distribution) at prophase vs. the total distance traveled, the average duration, and the average instantaneous speed of each kinetochore until it reaches the metaphase plate. The R^2^ value is based on linear regression of the scattered data and indicates no significant correlation.

Longitudinal tracking also enables us to test additional hypotheses about the search and capture process of kinetochores by microtubules. For example, a recent study suggested that the location of a kinetochore within the interphase nucleus can influence its propensity to missegregate during mitosis. It was reported via manually tracking kinetochores within a 2D plane, at 1 minute intervals, that kinetochores located near the periphery of the nucleus at the onset of prophase take longer to arrive at the metaphase plate and thus are more susceptible to erroneous microtubule attachments and mitotic errors^33^. We examined this question in our datasets by longitudinally tracking kinetochores in 3D at three or six second time intervals to accurately capture full kinetochore trajectories. Even with this high-temporal sampling, kinetochore tracking is not perfect due to difficulties in detection when kinetochores become too close in 3D or have poor signal to noise due to variability in protein expression level or photobleaching. We quantified the ratio of tracked kinetochores that could be unambiguously tracked forward in time from prophase or backward in time from anaphase onset and plotted the survival probability (i.e. the fraction of kinetochore trajectories that can be fully traced forward or backward from the starting point). The greatest difficulty in tracking occurs during prophase when kinetochores are clustered and display rapid intermittent motion during the initial stages of capture by microtubules. In our datasets, approximately 20 percent of kinetochores can be unambiguously tracked from the start of prophase through to anaphase onset (**Fig. S7**). Focusing specifically on these kinetochores, we tracked 899 trajectories from 33 cells total and plotted the duration, distance traveled, and average speed for each kinetochore as a function of its position within the prophase nucleus. Interestingly, we find no clear correlation between any of these values and the position of the kinetochore in the prophase nucleus (**Fig. 5 E-G**), although it’s not immediately clear if this might be due to differences in methodologies between our approach and prior studies.

## Discussion

In this work, we combine AI-based computer vision with LLSM to autonomously image rare cell events within a population. We demonstrate the utility of this approach to enable high content and high throughput imaging in both fixed and live-cell conditions. By scanning, localizing, and classifying hundreds of cells per second, smartLLSM dramatically exceeds the rate of manual sample search, and by automatically switching between inverted epifluorescent microscopy and LLSM, it both reduces the overall data burden and enriches the acquired data for the specific events of interest. Together, this allows a user to capture statistically significant numbers of 3D/4D datasets at rates that could not be accomplished via manual acquisition. This automated system allows quantification of both population-level statistics about mitotic stage as well as subtle effects of low-dose pharmacological perturbation to the sub-cellular localization of mitotic structures and proteins at specific mitotic stages. Automated sample search, live imaging, and longitudinal 4D kinetochore tracking allows identification of outlier kinetochores at different cell stages, reveals that certain mitotic stages are more prone to outlier events that others and explores the effects of drug perturbations on this process.

SmartLLSM builds upon recent work in automated, adaptive, or “self-driving” microscopes^17–19,34,35^. In particular, automated sample search and cell classification has been demonstrated previously using object segmentation, feature vector computation, and support vector machines (SVM) on commercially available confocal microscopes^17^. Timing data was not provided in these demonstrations, thus preventing a direct comparison to this work; however, due to the need to first segment cells and then compute feature vectors for each cell independently, our initial trials with SVM-based approaches were over an order of magnitude slower than our implementation with YOLOv5 on the same data. For example, generating the segmentation masks via Cellpose, the first step of our SVM classification test, took roughly 20-fold longer than the entire YOLOv5 pipeline. Although this could be explained in part by the specifics of how these processes were implemented, fully convolutional approaches are well-established to be faster than methods with separate detection and classification steps^21^. Combined with the fact that LLSM can be 1 to 2 orders of magnitude faster than point scanning confocal^6^, we estimate that our approach is between 10X - 100X faster than the prior state of the art, while achieving comparable accuracy for automated sample search and 3D imaging.

While our initial application focused on mitosis, our pipeline can be adapted to any biological phenomena that is morphologically distinct or that contains some unique combination of image properties. Potential examples include avoiding overexpression artifacts by automatically imaging fluorescent reporter cells whose protein expression levels fall within a certain range or identifying structures that form from heterogeneous cell-cell interactions such as T cell – cancer cell immune synapse formation^36,37^. However, a limitation for any supervised learning method, including smartLLSM, is that the network may behave unpredictably when presented with data that is outside the training dataset. As such, care must be taken to retrain the network if experimental conditions or the desired cellular events to be captured change. However, we anticipate that, for a given microscope and/or trained network, transfer learning could be used to dramatically cut down on the amount of new training data that might be necessary to adapt the instrument for a given task. Additionally, the network will still be subject to user bias in the training data annotation. As such, there is a risk that by only imaging cells that conform to preconceived ideals, the user may miss cells with unexpected and potentially interesting phenotypes. Finally, the switching time between inverted epifluorescent and LLSM modes on the current instrument is approximately 3 seconds which, without additional hardware upgrades, may be too slow to capture very rapid events like action potential-triggered neurotransmitter release^19^.

Overall, we demonstrate that smartLLSM bridges spatial and temporal scales, enabling mm-scale search and surveillance of cell populations and high-resolution 4D imaging of cellular dynamics. To facilitate the adoption of this approach and aid its adaptation to other types of microscopes, we provide our open-source annotated dataset, the annotation GUI, and the trained network. We anticipate that this automated approach will increase statistical power for observations of rare cellular events, increase experimental throughput by capturing relevant data at higher speeds than can be achieved via manual sample acquisition, and facilitate the screening of new pharmacological perturbations on a subset of rare cells or cell stages within heterogeneous populations.

## Methods

### Neural Network Training

#### i) Segmentation

To train our network, we utilized an initial data set of z-stacks of fixed cells expressing H2B-mScarlet captured through the inverted objective with epi illumination on the lattice light sheet instrument. For a given location, 41 z-planes were acquired at 1 μm spacing and 10 ms exposure to image a full volume above and below the coverslip. The field of view for each z-stack was 800 x 800 pixels in size (80 x 80 μm) and contained approximately 8 cells on average. The stages were then scanned to acquire z-stacks at 23,100 separate positions across the sample. The most in-focus position for each stack was evaluated based on the position with a minimum Shannon Entropy and was used for further processing. This process was repeated for three separate biological replicates. Segmentation of individual cells in each image was performed using Cellpose^25^, a deep learning-based segmentation algorithm. Batches of images were passed to the pretrained Cellpose network. The algorithm efficiently segmented out potential cells, allowing masks and cell outlines to be saved for 69,300 individual images. This allowed us to pass individual cells to our cell annotator and classification networks in subsequent steps. Of note, while Cellpose was able to segment clearly defined cells, it also picked up on blurry cells that lie in different focal planes as well as other non-cellular debris that was picked up while imaging. We utilized general image features (e.g., mask diameter) along with classification categories while annotating (to be learned by the classification network) to account for these blurry cells and erroneous Cellpose masks.

#### ii) Classification

In order to generate labeled training data from the raw images (i.e., label the mitotic stage of each cell), we developed a custom interface in Python (**Fig. S8**). The main GUI consisted of a split view, with the right half displaying the current cell to be labeled and the left half displaying the entire image (with the current cell outlined) to provide local context for the user. This was especially useful when classifying cells in stages such as late anaphase and telophase. In these cases, Cellpose tended to segment the single splitting cell into two separate cells. However, using the global context, it became easy to classify these cells as the second daughter cell could be seen close by the current cell. Moreover, we were able to fine tune our annotation options to account for any artifacts in the Cellpose masks (e.g., debris or cells that were clipped by the edge of an image).

We utilized an iterative approach to bootstrap and expedite annotation by embedding a neural network within the annotation GUI to “pre-screen” images that contained mitotic cells. Because mitotic cells represent a small fraction of all cells in the images, the class distribution is highly skewed and many images contained only interphase cells. Therefore, we first manually scanned the data to generate a small subset of curated cells (e.g. 10 cells for each class). We used these images to train a three-category neural network classifier that could discriminate between interphase, blurry, and mitotic cells (**Fig. S1**). Before being passed to the network, we cropped out a 286 x 286 pixel region centered each cell. Other cells present off-center, but still in the crop, were retained as opposed to masking each individual cell since it seemed to improve classification results. Next, data augmentation was performed using TensorFlow’s “ImageDataGeneration” function. This function preprocessed and augmented the cells via sample-wise standardization and image rotation, flipping, and shearing. The trained network was then implemented in the cell annotator, allowing each image to be “pre-scanned” to determine if the image contained only interphase cells or if cells in any mitotic class were present. This process could be refined by changing the threshold value for classifying cells as mitotic, allowing for a trade-off between the likelihood of images presented to the user containing mitotic cells vs. the likelihood of erroneously discarding useful images to annotate. This allowed for a much more efficient method to acquire training data. This “in-line” neural network was periodically retrained as we acquired more annotated data allowing for more efficient filtering of images and ultimately a more accurate classifier.

#### iii) YOLO Implementation (smartLLSM Neural Network)

Once we obtained the training data, we utilized the state-of-the-art, open-source YOLO architecture^21^ to rapidly locate and classify cells within the microscope images into the following categories: interphase, prophase, prometaphase, metaphase, anaphase, telophase, and blurry. YOLO is a single-shot detector (it does not have a separate region proposal step), prioritizing detection speed while still maintaining high classification accuracy. We refer the reader to the well-maintained and documented YOLOv5 repository for detailed instructions for training YOLO on custom datasets^38^. In short, we transformed our labeled data to a YOLO-compatible format with uniform square bounding boxes (286×286, to match our annotation procedure). YOLO is trained on the entire microscope field of view (versus individual cell crops, as per our annotation procedure). As such, we split the 5659 annotated images into a training (70%, 3981 images) and test set (30%, 1678 images). In the training set, 33879 total cells were used: 84 anaphase, 5787 blurry, 26284 interphase, 631 metaphase, 293 prometaphase, 258 prophase, and 542 telophase. In the test set, 15154 total cells were used: 29 anaphase, 2814 blurry, 11592 interphase, 260 metaphase, 141 prometaphase, 132 prophase, and 186 telophase.

We trained the YOLOv5s (“small”) model using default settings for 195 epochs (early stopping, best model used) in 2.8 hours using a Linux workstation equipped with an NVIDIA GeForce RTX 2080 Ti GPU with 12 GB VRAM. Network performance results are shown in **Figs. 1B,1C, S8-S10**. To validate the localization accuracy of YOLO, we compared the center of YOLOs bounding boxes to the center of the original Cellpose masks and found good agreement between YOLO and Cellpose (**Fig. S9**) which was sufficient for accurately centering the cells for 3D LLSM imaging.

### Microscopy optical path

The optical path for smartLLSM is based on a modified version of the instrument described in Chen et al.^6^. Key modifications relevant to this work are the use of a 0.6 numerical aperture (NA) excitation lens (Thorlabs, TL20X-MPL), and 1.0 NA detection lens (Zeiss, Objective W “Plan-Apochromat” x20/1.0, model # 421452-9800), and a matching 1.0 NA objective lens located below the sample to allow for high-resolution epifluorescent inverted imaging.

### smartLLSM operation

Microscope control is accomplished with a custom developed and freely available LABVIEW software^6^. For inverted imaging, we set the ROI of the camera to 800 by 800 pixels and the scanned a tiled array of 10 by 10 positions (in total 800 μm by 800 μm). Inverted imaging was done with an exposure time of 50 ms resulting in approximately 6 s to cover the 800 μm by 800 μm regionwhen including stage settling time. To scan the entire coverglass, we set a list of positions in a 19 by 6 grid covering an area of 16 mm by 6 mm (**Fig. 2A**). To compensate for the any coverglass warping over this large area, we manually tuned the z-focal position in a 3 by 3 grid covering the same area and then fit these positions to a two dimensional second order polynomial to program the z-focal position of each point in the 19 by 6 grid. This 3D position list is then used to perform “Sample Finding” scanning to generate the input data for the deep learning network.

The detection of mitotic cells is done by a Python script that runs in parallel to the LABVIEW instrument control software. Images acquired by LABVIEW are passed to the Python script to be processed by the trained YOLOv5 network. After processing, the Python script outputs a list of coordinates of mitotic cells together with the classification probability from YOLOv5 (**Fig. S10**). We applied a threshold at 0.2 on the classification likelihood to filter out false positive detection of mitotic cells. The threshold was determined based on manual evaluation of the network performance and the same threshold was applied to both live cell and fixed cell imaging. For cells in anaphase or telophase, the YOLOv5 network tends to independently localize each of the separated daughter cells. To account for this, we averaged any localizations from these classes that occurred within 10 μm of each other to accurately identify the center of the dividing cell (**Fig. S10 d-g**). Once a targeted position is generated from the Python script, its coordinates are passed to the Labview software to move the stage to the target position and switch the microscope to LLSM mode. We performed LLSM scanning by moving the sample stage laterally, in the plane of the coverslip, with an “x” step-size of 400 nm (this results in 215 nm translation along the detection optical axis due to an angle of 32.5 degrees between detection focal plan and sample coverglass) over a range of 60 μm. For live-cell imaging, we used a 10 ms exposure time leading to a sampling rate of 2.8 s per volume for the first 100 frames, which typically covers prophase and early prometaphase, and a 20 ms exposure time leading to a sampling rate of 5.8 s per volume for the rest 450 frames, which cover the rest of the mitotic process (in total approximately 50 minutes per cell). This maximizes the sampling rates during prophase to capture the fast-moving kinetochores while minimizing photobleaching and phototoxicity. For fixed cell imaging where rapid acquisition was less critical, we increased the exposure time to 50 ms and utilized sequential exposures for each wavelength to minimize spectral bleedthrough. This resulted in a volumetric sampling time of 22 s per cell. Detailed experimental parameters for all datasets are provided in **Table S2**.

### RPE cell line generation, cell culture, and sample preparation

All cells used in this study were maintained in hTERT RPE-1 growth media consisting of DMEM/F12 (Thermo Fisher Scientific, 11320033) media supplemented with 10% FBS (Omega Scientific FB-11), 100 U/mL Penicillin-Streptomycin (Thermo Fisher Scientific, 15140122), and 0.01 mg/mL hygromycin B (Thermo Fisher Scientific, 10687010) unless otherwise noted. The cells were cultured in 25cm 2 or 75cm 2 dishes without a fibronectin coating. hTERT RPE-1-mScarlet-H2b+mNeon-CENPA cells were generated from parental lines (RRID: CVCL_4388, ATCC) using piggyBac transposon-based methods^39^. The cDNA for mScarlet-H2b and mNeon-CENPA genes was synthesized by GeneScript, then ligated into piggyBac plasmids that conferred resistance to either Blasticidin or Gentymycin^40,41^. 0.3 x 10^6^ hTERT RPE-1 cells were electroporated with 4 μg of mScarlet-H2b, 4 μg of mNeon-CENPA, and 2 μg of piggyBac Transposase (Systems Biosciences, PB210PA-1), using the Neon Transfection system. Immediately following electroporation, cells were plated in 35mm dishes with antibiotic-free hTERT RPE-1 growth media. The media was replaced with fresh antibiotic-free media 6 hours after plating and again the following day. After 48 hours to allow the cells to recover from electroporation, cells were maintained in selection media containing 800 μg/mL of G418 Sulfate (Thermo Fisher Scientific, 10131027) and 6 μg/mL of Blasticidin (Goldbio, B-800-25) for two weeks. The hTERT RPE-1-mScarlet-H2b+mNeon-CENPA cells were then expanded in hTERT RPE-1 growth media and harvested for cryopreservation in a mixture of 40% DMEM/F12, 60% FBS, and 10% DMSO (Millipore-Sigma, D2650-100ML). hTERT RPE-1-mScarlet-H2b+mNeon-CENPA cells were used for experiments for up to 20 passages after thawing.

For imaging, 25 mm #2 coverslips (Warner Instruments, 640722) were sonicated in 1M KOH (Millipore-Sigma 484016-1KG) for 30 minutes, washed with diH2O then sonicated in diH2O for 30 minutes. Immediately after sonication in diH2O, cover slips were dried with compressed nitrogen then stored a plastic dish lined with lens paper for a maximum of three weeks. One day prior to imaging experiments, cleaned coverslips were incubated with 10 μg/mL of fibronectin (Stemcell Technologies, 07159)) at 37°C for 30 minutes. Immediately thereafter, hTERT RPE-1-mScarlet-H2b+mNeon-CENPA cells were plated on fibronectin coated coverslips at a density of 3.4 x 10^4^ cells/cm^2^.

### Taxol Treatment

Two days prior to imaging experiments, hTERT-RPE-1-mScarlet-H2b+mNeon-CENPA cells were plated on fibronectin coated coverslips at a density of 1.7 x 10^4^ cells/ cm^2^. One day prior to imaging experiments, media was replaced with hTERT RPE-1 growth media supplemented with 5 nM Taxol (Millipore-Sigma, PHL89806-10MG). Cells were maintained in Taxol supplemented media for 20 hours. For live-cell experiments, cells were imaged in FluoroBrite (Thermo Fisher Scientific), A1896701) supplemented with 10% FBS, 100 U/mL Penicillin-Streptomycin and 5 nM Taxol. For fixed cell experiments, cells were washed once with 37°C PBS and immediately fixed in a 4% paraformaldehyde-PBS (Thermo Fisher Scientific, 50-980-487) solution for 12 minutes at room temperature. Cells were washed once with PBS for 10 seconds followed by 3 washes with PBS for 5 minutes. Following the final wash step, cells were stored for a maximum of 72 hours at 4° C prior to imaging.

### Immunofluorescence

hTERT RPE-1-mScarlet-H2b+mNeon-CENPA cells were plated on fibronectin-coated dishes at the specified cell densities and times above prior to immunostaining. The cells were washed once with 37°C PBS and immediately fixed in a 4% paraformaldehyde-PBS solution for 12 minutes at room temperature. The cells were then washed with PBS and permeabilized with a 0.5% Triton-PBS (VWR, 0694-1L) solution for 20 minutes. The cells were washed with PBS and blocked in a normal goat serum-PBS (NGS-PBS) (Thermo Fisher Scientific, ICN642921) solution for 1 hour. The cells were then incubated in a 1:250 Anti-β-Tubulin mouse monoclonal antibody (Millipore-Sigma, T5293-.2ML)-NGS-PBS solution for 2 hours. The cells were washed with PBS and incubated with a 1:500 solution of Alexa Fluor 647 (Thermo Fisher Scientific, A-21245) conjugated goat anti-mouse secondary antibody NGS-PBS solution for 2 hours. Cells were washed with PBS and stored for a maximum of 48 hours at 4° C prior to imaging. To minimize the loss of less-adherent mitotic cells, all incubation steps were performed without rocking and all aspiration steps were performed using a pipette instead of a vacuum. The protocol for all PBS wash steps, except for the initial washing prior to fixation, consisted of 1 wash with PBS for 10 seconds followed by 3 washes with PBS for 5 minutes. All steps were performed at room temperature unless otherwise noted.

### Cell synchronization

Four days prior to imaging, hTERT-RPE-1-mScarlet-H2b + mNeon-CENPA cells were plated on fibronectin coated coverslips at a density of 6000 cells/ cm^2^ for thymidine (Sigma, T9250) blocked samples and 3000 cells/cm^2^ for control samples. On day 2 after plating, media for the thymidine blocked samples was replaced with 8 nM thymidine supplemented media. Control cells received fresh media. Cells were allowed to incubate for 18 hours. After 18 hours, all samples were washed 3 times with warm media and received fresh media. Cells were allowed to incubate for 8 hours. Media for the thymidine blocked samples was then replaced with 8 nM thymidine supplemented media and control cells received fresh media. Cells were again allowed to incubate for 18 hours. After 18 hours, all samples were washed 3 times with warm media and received fresh growth media. Cells were then allowed to incubate for 5 hours before being moved to the microscope for imaging.

### Imaging processing, kinetochore tracking and trajectory quantification

Because of the angle between the detection focal plane and the sample plane, raw lattice light sheet images are first deskewed^6^. The deskewed images are then deconvolved with the corresponding experimentally measured point spread function using Richard-Lucy method for 10 iterations^42^.

#### i) Kinetochore localization and tracking

For kinetochore localization, we applied a difference of Gaussian filter to the 3D deconvolved images of the kinetochore channel and then fit the local maxima of each spot with a 3D Gaussian function using a maximum likelihood estimator (MLE). To track kinetochores across time points, we utilized a modified Matlab code based on uTrack^31^ with a maximum search radius of 10 pixels (~ 1 μm). To remove spurious localizations, we discarded any tracks with a length less than 10 frames. To identify the cell stage from the live-cell time lapse movies of kinetochore trajectories, we manually determined the first frame corresponding to the onset of mitotic stage based on kinetochore motion and organization. We defined the “contracting stage” as the starting point where kinetochores move rapidly inward after prophase, prometaphase as the endpoint of this contracting process, metaphase as the time point where all kinetochores aligned to the metaphase plate and anaphase as the time where kinetochores start to move outward toward the two daughter cells (**Movie S3**).

#### ii) Quantification of kinetochore spatial distribution and metaphase plate orientation

To compensate for varying cellular orientation and to quantify changes in the kinetochore arrangement under different conditions, we first applied principal component analysis of the localizations of kinetochores in each image using the PCA function in Matlab (The Mathworks). This function extracts the eigenvalues, the eigenvectors, and the projected coordinates of the localizations to the eigenvectors of the data. The orientation of the metaphase plate (θ, **Fig. 3D, E**) is defined as the angle between the normal vector of the sample plane (in our case the z direction) and the eigenvector associated with the smallest eigenvalue. These values provide a normalized metaphase plate reference frame (defined as the projected coordinates along the first and second eigenvectors) that can be used to register other channels within the same dataset. The normalized variation of kinetochore coordinates along the first and second eigenvectors respectively can be used to define an affine transform, 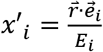, where 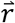 is the kinetochore location in the camera coordinate, and 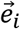, is the i^th^ eigenvector and *E_i_*, is the i^th^ eigenvalue. This transform operates on either the 3D point cloud of kinetochore locations or on the pixel coordinates within the images themselves. This affine transform can then be used to rotate and stretch the images of chromosomes, microtubules, and kinetochores to register them into the same coordinate frame and account to variations in cellular orientation and size. We generated heatmap images of the distributions for kinetochores, chromosomes, and microtubules based on the histogram of positions (kinetochores) or immunofluorescence intensity (chromosomes and microtubules) aggregated across all cells in each condition (**Fig. 3F**).

We quantified the anisotropy of the kinetochore distribution (**Fig. 3H**) as 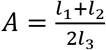, where *l*_1_, *l*_2_, and *l*_3_ are the largest to smallest ordered eigenvalues respectively. To quantify the orientation of the metaphase plate in live-cell movies, we plotted the metaphase plate orientation (defined above) and as a function of time after the start of prometaphase (**Fig. 4D**). We then fit the metaphase plate orientation vs. time curve from each cell to a sigmoid function, which follows the form of 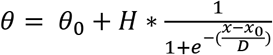**(Fig S5)**. From these fits, we extracted the total reorientation angle H and the time at which the reorientation finishes (defined as *x*_0_ + *D*).

#### iii) Calculation of kinetochore speed

To compensate for the uneven sampling rate during prophase and the rest of the mitotic process (2.8 s per volume for the first 100 frames and 5.8 s per volume for the last 450 frames), we first interpolated the kinetochore tracks over the first 100 frames using a sampling rate of 5.8 s per volume. To compensate for global cell motion or microscope drift, we subtracted the average translational motion of all kinetochores between frames. We then calculated kinetochore speed as the distance a kinetochore traveled between two neighboring interpolated frames.

To segment the speed distribution at different mitotic stages, we manually annotated the onset of contracting, prometaphase, metaphase and anaphase in each mitotic cell time-lapse as described above. For the non-splitting cells treated with 1 nM or 5 nM Taxol, there is no anaphase onset time point. To generate probability distributions of the kinetochore speed, we aggregated the computed speed at each time point in a given mitotic stage of all kinetochores across all cells. To generate the box plot of medians of the velocity distribution, we calculated the median velocity of all kinetochores within each cell across all time points for a given mitotic stage.

#### iv) Abnormal kinetochore detection

Because there can occasionally be extra cells present within a live-cell movie in addition to the mitotic cell of interest, we first identified the kinetochores corresponding only to the mitotic cell of interest by applying the DBSCAN function in Matlab with a search radius of 50 pixels (5.4 μm) and a minimum cluster size of 50 points. Localizations within the largest cluster correspond to the dividing cell of interest and are utilized for further analysis. To account for spurious localizations that may results from camera noise or other sources, we additionally filtered localizations to only include tracks with a minimum length of 10 frames. We then project the remaining kinetochore localizations along the eigenvectors based on PCA analysis described above, and applied “rmoutliers” function in Matlab to detect abnormal kinetochores. We used “gesd” method (generalized extreme studentized deviate test) with a detection threshold of 0.05. For time points after the anaphase onset, we detect the two daughter cells using the DBSCAN function in matlab with a search radius of 40 pixels (4.3 μm) and a minimum cluster size of 40 points and then detect outliers for each cluster using the same “rmoutliers” function as described above. The probability for observing an outlier kinetochore at a given stage is calculated as 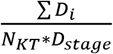, where *D*_i_ indicates the duration of an outlier over which kinetochore i was considered an outlier, *N_KT_* is the total number of kinetochores in a cell and *D_stage_* is the duration of a specific mitotic stage.

#### v) Analysis of longitudinal kinetochore tracks

To analyze the relationship between kinetochore location during prophase and kinetochore dynamics and location in the metaphase plate, we only included kinetochores with trajectories that span all time points from prophase to the onset of metaphase. We calculated the Mahalanobis distance of each kinetochore to the group centroid and normalized it to a value between 0 and 1 (0 for the kinetochore at the center and 1 for the kinetochore at the edge). To calculate the time point where the kinetochore reaches the metaphase plate, we generate the convex hull that covers the kinetochore distribution during metaphase, and detected the first time point when the kinetochore is included in the convex hull. We then calculated the traveled contour length of the kinetochore trajectory and the median instantaneous speed throughout each kinetochore track.

### Statistics and reproducibility

No statistical method was used to predetermine the sample size. Data sets for fixed cells were repeated for three independent biological replicates each with two cover glass samples for each condition. Data for mitotic wave detection was repeated for two biological replicates with one coverslip for each replicate and condition. Datasets for live cell movies comparing control and Taxol conditions were repeated for three independent biological replicates for each condition.

For fixed samples, we used a T-test to compare the proportion of cells in each mitotic stage between control and Taxol treated conditions and a chi-square test of independence to determine whether there is a change in the overall distribution of mitotic stages between control and Taxol conditions (**Fig. 3B**). We used a Kolmogorov–Smirnov test to determine if there are differences in the distribution of the metaphase plate orientation and kinetochore anisotropy between control and Taxol conditions (**Fig. 3E, H**).

For live-cell movies, we used a T-test to compare if there is a significant difference in the median kinetochore speed between control and various Taxol-treated conditions and to compare if there is a significant difference in the parameters extracted from sigmoid fits of the metaphase orientation curves (**Fig. 4 C, Fig. S5**). We used a Kolmogorov–Smirnov test to determine if there is a significant difference between the probability, duration, and normalized probability of abnormal kinetochores in different mitotic stages (**Fig. 5B-D**). We used linear regression to test if there is a linear relationship between a kinetochore’s position in prophase and the traveled distance, the duration, and the speed of the kinetochore between prophase and metaphase (**Fig. 5E-G**).

## Data Availability

Due to the inordinate size of the image data (~50TB), it is not currently feasible to deposit this into a central repository; however, all datasets underlying the results in this manuscript are available from the corresponding author upon request. To the extent possible, the authors will try to meet all requests for data sharing within 2 weeks from the original request.

## Code Availability

The source code, annotation GUI, the library of annotated training data, and the trained YOLOv5 network generated in the current study are available at: https://github.com/nel-lab/smartLLSM. Code is provided under The MIT License for open source software, a permissive license approved by the Open Source Initiative. Specific terms can be found here: https://opensource.org/licenses/MIT.

## Acknowledgements

We thank Katelyn Heath and Michaela Clynes for assistance with annotating images. We thank Dr. Gokul Upadhyayula for assistance with the single particle tracking code and Dr. Tarun Kapoor, Dr. Michael Emanuele, and Dr. Adam Palmer for helpful discussions and feedback on the manuscript. This work was funded in part by grants from the National Institutes of Health (1DP2GM136653) awarded to W.R.L..

W.R.L. acknowledges additional support from the Searle Scholars program, the Beckman Young Investigator Program, and the Packard Fellowship for Science and Engineering.

## Contributions

W.R.L. conceived the project. J.T., Y.S., and A.G. generated the annotation software and trained YOLO network. D.E.M assisted with integrating the YOLO network together with the microscope control software. T.A.D and C.Y. assisted with sample preparation. Y.S. and W.R.L performed the imaging experiments, analyzed data, and wrote the manuscript with feedback from all authors. W.R.L. supervised and directed the project.

