## Supplementary Information for "Smart Lattice Light Sheet Microscopy for imaging rare and complex cellular events"

**List of SI movies**

Movie 1: Fixed-cell gallery of 3D datasets for different mitotic stages showing CENPA (magenta), H2B (cyan) and tubulin (yellow)

Movie 2: Two-color live-cell mitosis movie showing CENPA (magenta) and H2B (cyan)

Movie 3: Maximum intensity projection of a high-speed volumetric mitosis time-lapse showing CENPA and overlaid kinetochore tracks. For display purposes and to correct for photobleaching, each frame of the movie was linearly rescaled between [0 0.5*max_frame] where max_frame is the maximal pixel value at a given time point.

Movie 4: Maximum intensity projections of kinetochore dynamics in live cell movies showing control and 5 nM Taxol treated conditions. For display purposes and to correct for photobleaching, each frame of within the control and Taxol-treated datasets was linearly rescaled between [0 0.5*max_frame] where max_frame is the maximal pixel value for a given condition at a given time point.

Movie 5: Gallery of kinetochore dynamics in live-cells showing outlier kinetochore classification and tracking. For display purposes and to correct for photobleaching, each frame of each cell within the movie was linearly rescaled between [0 0.5*max_frame] where max_frame is the maximal pixel value at a given time point.

### Kinetochore + Chromosome

| Condition | Mitosis finish rate | Number of cells |
|---|---|---|
| 50 ms exp, 0.8 prct, 100 um range, 66 s period | 83% | 6 |
| 50 ms exp, 0.8 prct, 100 um range, 32 s period | 100% | 3 |
| 50 ms exp, 0.8 prct, 100 um range, 22 s period | 100% | 3 |
| 50 ms exp, 0.8 prct, 100 um range, 16 s period | 50% | 4 |

### Kinetochore only

| Condition | Mitosis finish rate | Number of cells |
|---|---|---|
| 50 ms exp, 1.2 prct, 100 um range, 15 s period | 100.00% | 8 |
| 20 ms exp, 3 prct, 100 um range, 9 s period | 100.00% | 1 |
| 10 ms exp, 6 prct , 100 um range, 6 s period | 84.62% | 13 |
| 5ms exp, 12 prct, 100 um range, 2 s period* | 83.33% | 6 |
| 2ms exp, 30 prct , 100 um range, 1 s period* | 0.00% | 4 |

**Table S1: Parameter search to balance the signal-to-noise ratio (SNR) and phototoxicity for live-cell imaging.** The top table represents the mitotic success rate for cells simultaneously imaged for kinetochores and chromosomes under 488 nm and 560 nm excitation, and the bottom represents the mitotic success rate for cells imaged for kinetochores under 488 nm excitation only. * indicates cases with significant photobleaching in kinetochore signal at the end of imaging.

<table>
<tr><th>Condition</th><th>Figure</th><th>Illumination mode</th><th>Exposure</th><th>Laser λ, power (at pupil)</th><th>Detection filter</th><th>Scan range</th><th>Scan step</th><th>Volume rate (s)</th></tr>
<tr><td>Live cell, fast</td><td>Fig 4, Fig 5, Fig S3-S7</td><td>LLS (MB-square, NA 0.4/0.3)</td><td>10 ms</td><td>488 nm, 134 µW</td><td>525/50 bandpass filter</td><td>60 µm</td><td>400 nm</td><td>2.8 s</td></tr>
<tr><td>Live cell, slow</td><td>Fig 4, Fig 5, Fig S3-S7</td><td>LLS (MB-square, NA 0.4/0.3)</td><td>20 ms</td><td>488 nm, 109 µW</td><td>525/50 bandpass filter</td><td>60 µm</td><td>400 nm</td><td>5.8 s</td></tr>
<tr><td rowspan="3">Fixed cell</td><td rowspan="3">Fig 3</td><td rowspan="3">LLS (MB-square, NA 0.4/0.3)</td><td rowspan="3">50 ms</td><td>488 nm, 975 µW</td><td>525/50 bandpass filter</td><td rowspan="3">60 µm</td><td rowspan="3">400 nm</td><td rowspan="3">22 s</td></tr>
<tr><td>560 nm, 29 µW</td><td>561 long pass filter</td></tr>
<tr><td>642 nm, 240 µW</td><td>643 notch filter</td></tr>
<tr><td>Sample scan</td><td>Fig 2A (i)</td><td>Epi</td><td>50 ms</td><td>560 nm, 207 µW</td><td>561 long pass filter</td><td>800 µm by 800 µm</td><td>80 µm</td><td>6 s</td></tr>
</table>

**Table S2: Image parameters for smart LLSM operation.** "Live cell, fast", "Live cell, slow", and "Fixed cell" are the imaging parameters used for 3D volumetric scanning through lattice light sheet microscopy, and "Sample scan" is the imaging parameters used for sample finding through the inverted objective.

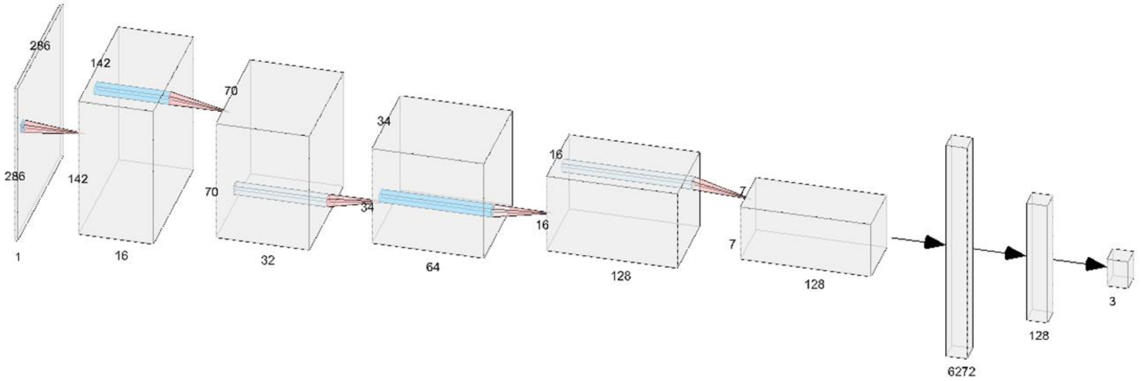


**Figure S1: A sketch of the architecture of the boot-strapping convolutional neural network used for data annotation.** Each segment is composed of five layers of 3x3 2D convolutional layers with relu as the activation function followed by a 2x2 max pool layer. The data are then passed on to a flattening layer, followed by two fully connected layers with relu activation function and a drop rate of 0.5. A fully connected layer with soft max activation function is used to yield the final 3-class result.

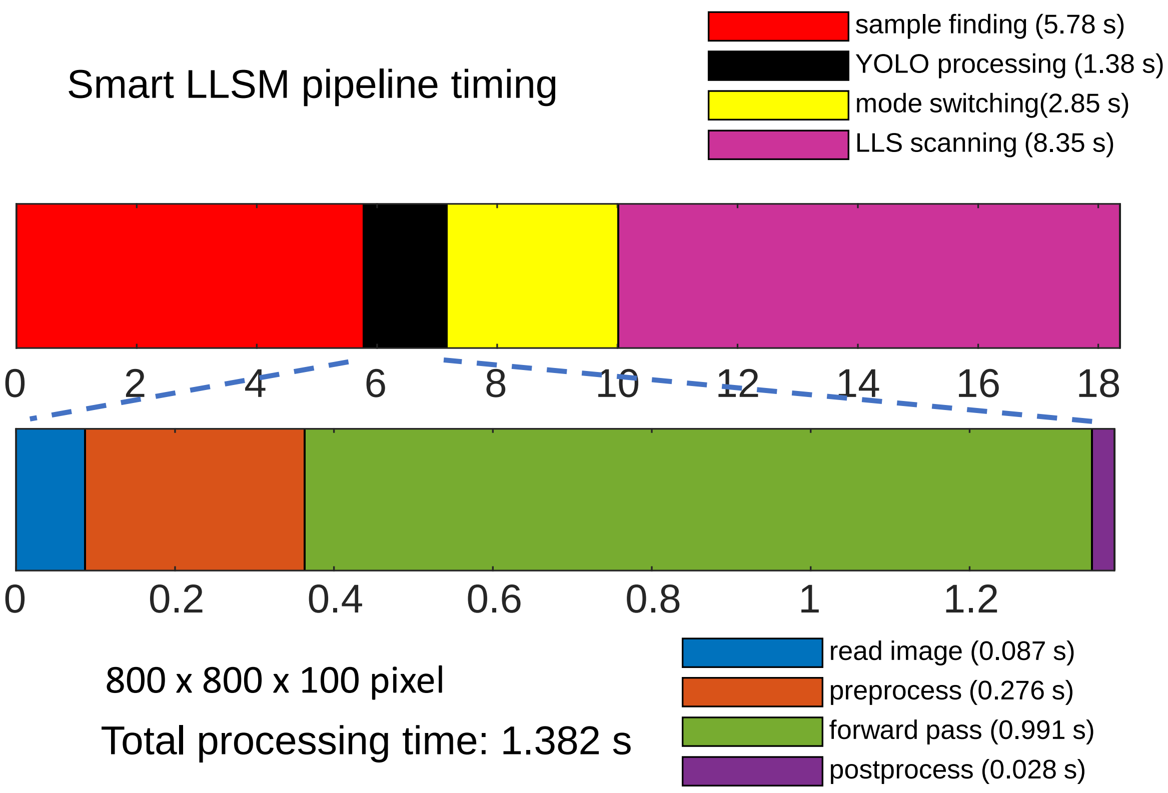


**Figure S2: smartLLSM timing.** The top row shows the overall timing diagram for smartLLSM operation. The sample finding is performed over a 10 x 10 grid (100 2D images) that covers an 800 µm by 800 µm region at 50 ms exposure time for each image. For this example, 3D LLS scanning is done in a single color at 50 ms exposure time covering a scan range of 60 µm with 0.4 µm step size. Live-cell imaging was done over a similar range, but with either 10 ms or 20 ms exposures. The bottom row is the detailed timing of YOLOv5 network processing.

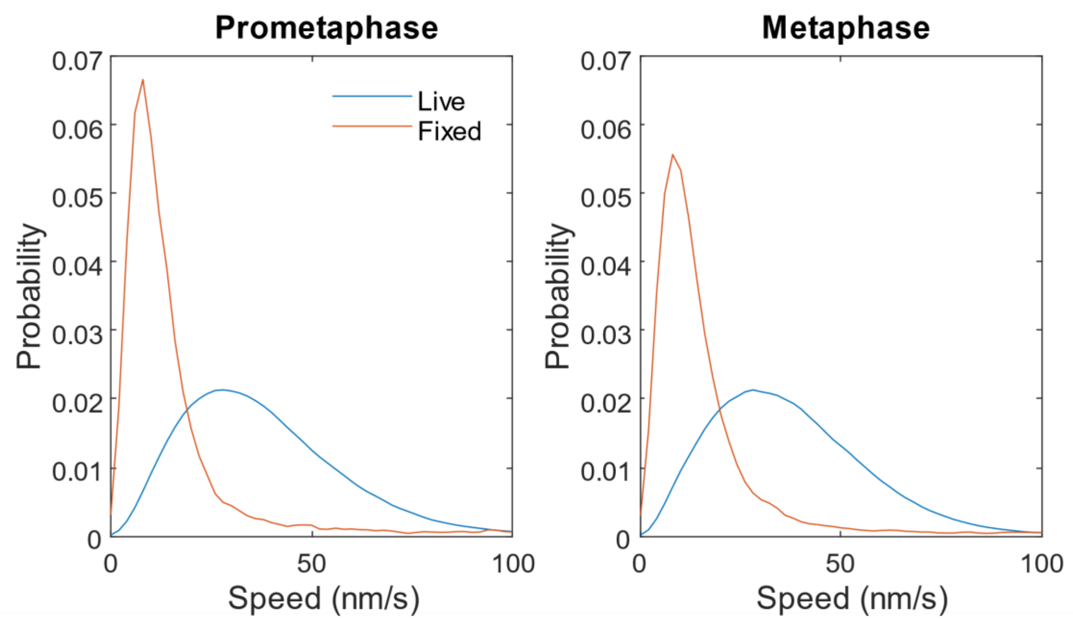


**Figure S3: Estimate of kinetochore tracking error.** Distribution of instantaneous speed in chemically fixed (orange) and live cells (blue) in prometaphase (left panel) and metaphase (right panel). The median kinetochore speed in fixed cell is 10.8 nm/s for prometaphase and 12.4 nm/s for metaphase. For live cells, this value is 35.2 nm/s and 35.1 nm/s, respectively.

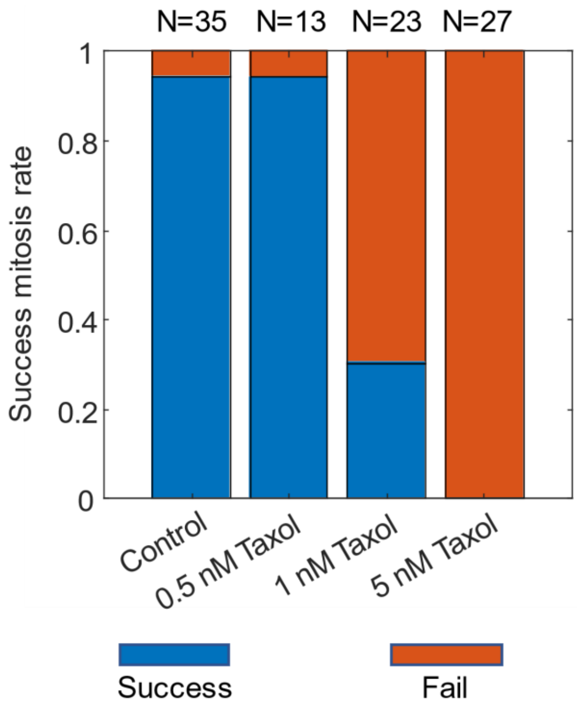


**Figure S4: Mitotic completion rate under different drug conditions.** Sample size on top indicates the total number of cells imaged for each drug concentration.

(a)

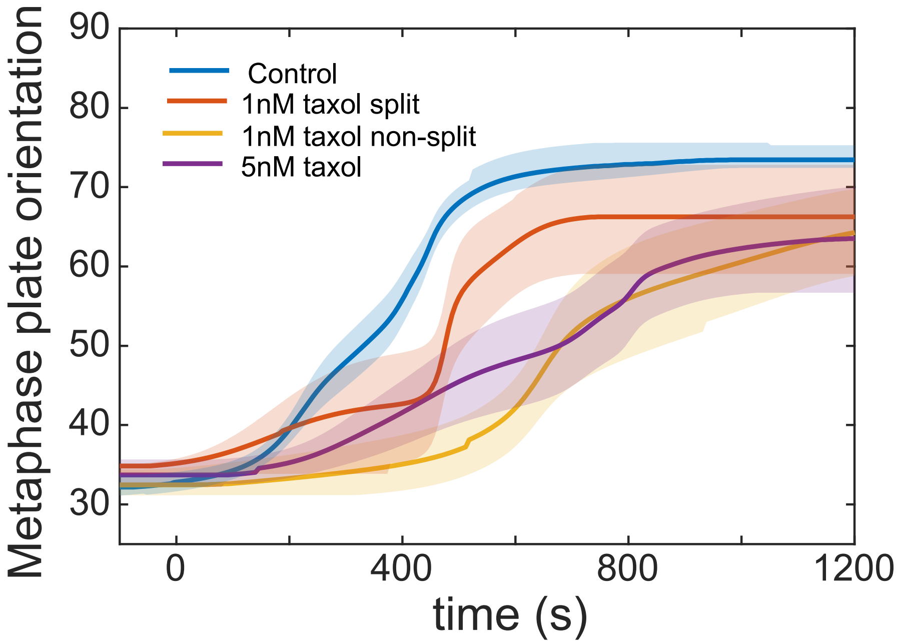


(b)

| | **1nm split** | **1nm nonsplit** | **5nm** |
|---|---|---|---|
| Angle change | 0.28 | 0.0555 | 0.0729 |
| Reach plateau | 0.04 | 2.6E-6 | 6.0E-6 |

**Figure S5: Sigmoid fitting of metaphase plate orientation in live-cell movies.** (a) Sigmoid fitting of metaphase plate orientation over time in each cell. Line indicates the mean and the shade indicates the standard error of the mean of the fitted curves. (b) Significance tests on the angle (height of the sigmoid function) and the time point when the metaphase plate orientation reaches a plateau (duration + offset). The table shows the p-value from the t-test comparing Taxol treated condition with control. See Methods for details on parameter fitting.

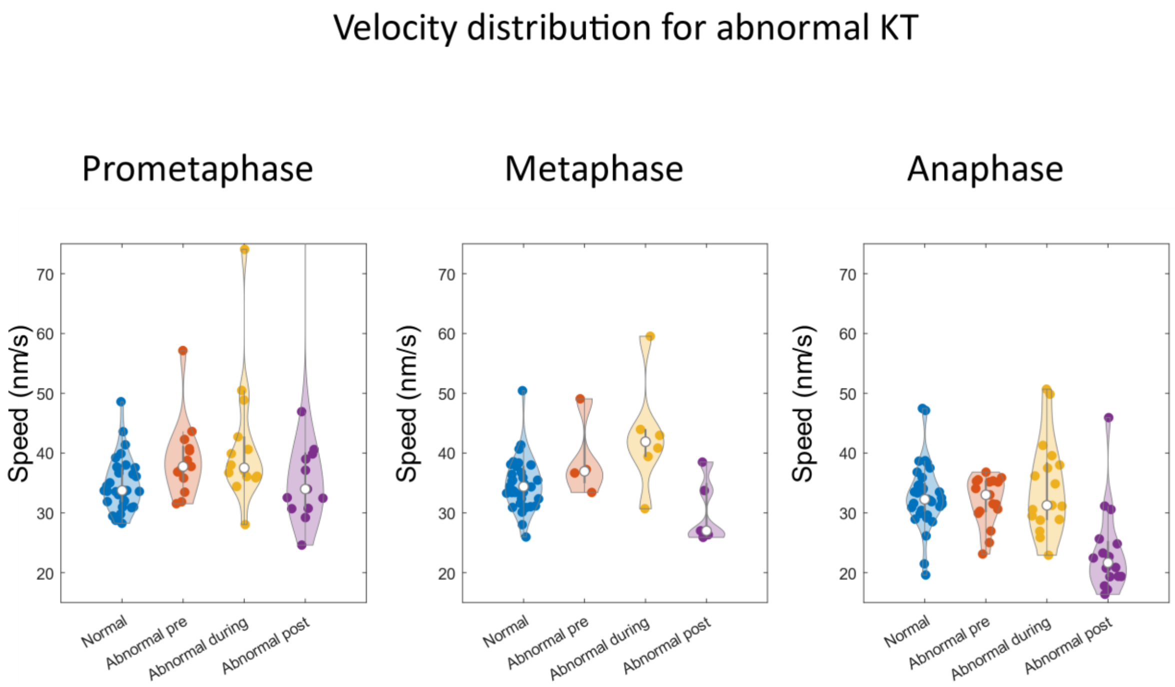


**Figure S6: Analysis of outlier kinetochore speed.** Violin plots of the speed of normal kinetochores and outlier kinetochores before, during and after being called as an outlier. The left panel shows prometaphase, the middle panel show metaphase, and the right panel shows anaphase.

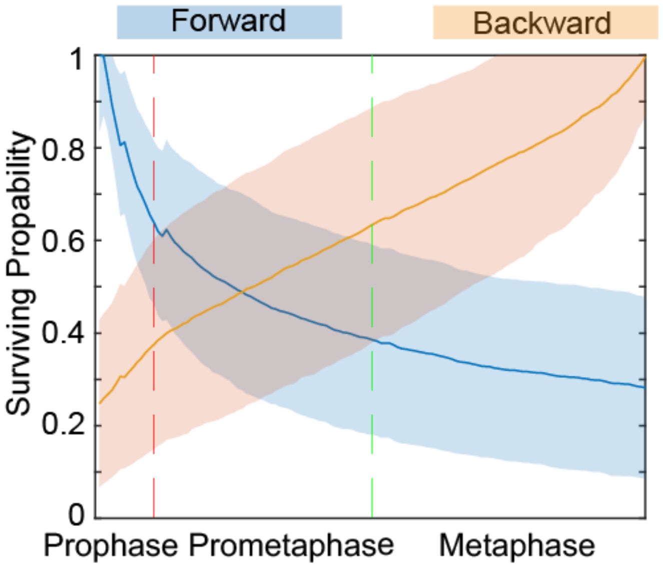


**Figure S7: Survival probabilities of tracked kinetochores throughout mitosis.** The survival probability is calculated as the proportion of kinetochores that can be continuously tracked from prophase (blue) forward or from anaphase (orange) backward in time. The solid line indicates the average surviving probability across cells and the shade indicates the standard deviation.

(a)

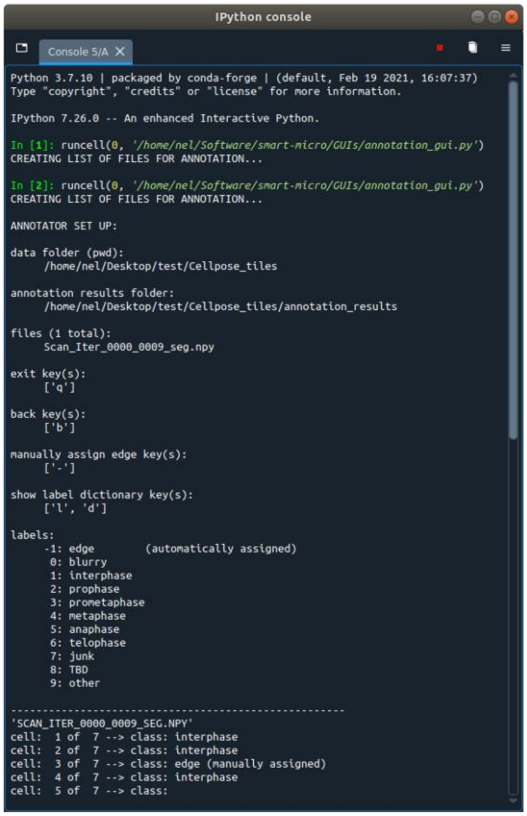


(b)

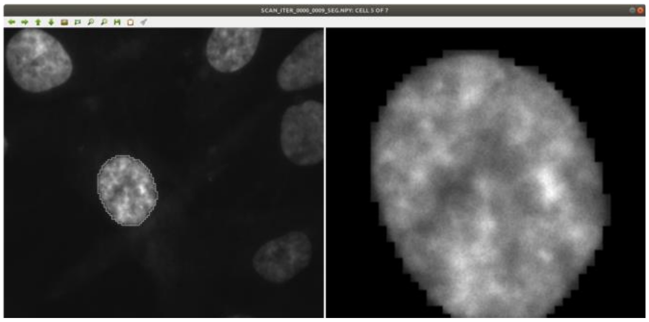


**Figure S8: GUI for smartLLSM annotation.** (a) Annotation GUI input interface. Features include the ability efficiently annotate cells, relabel previously annotated cells within a tile, and in-line documentation of annotation results. (b) Annotation GUI window. The split screen window displays the entire tile on the left half with the individual cell to be labeled denoted with a white border, allowing for global perspective when annotating cells. The cell to be annotated is displayed in a zoomed-in view on the right half.

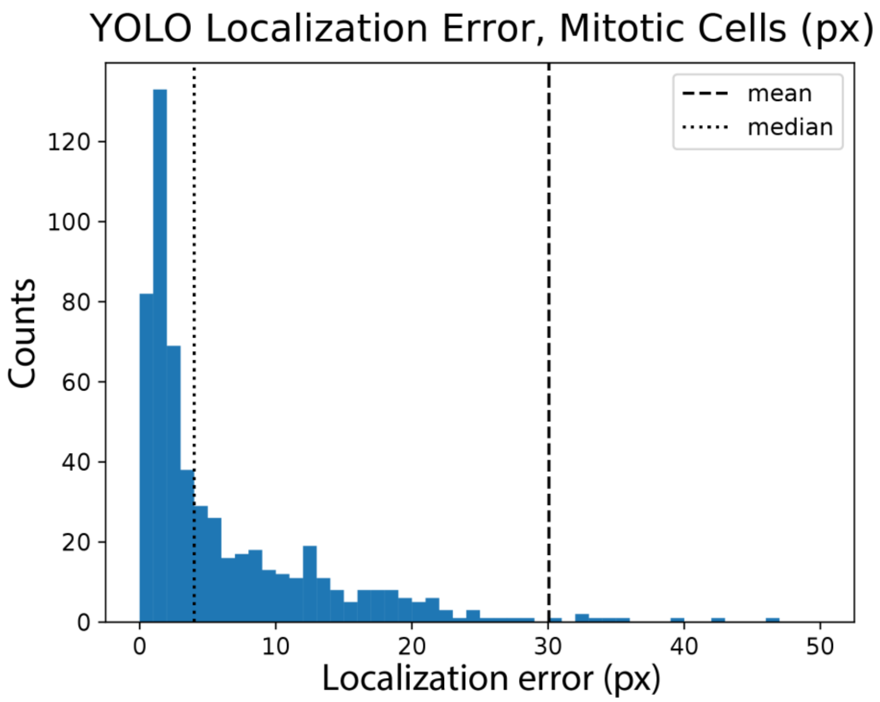


**Figure S9: YOLOv5 localization error.** Localization error in pixels was calculated between the center of mass of the Cellpose mask (ground truth) and YOLOv5 bounding box centroids from the same test set used evaluating YOLOv5 performance metrics. Erroneous Cellpose masks were filtered out based on size and classification. 642 mitotic cells were compared between Cellpose and YOLOv5 with a mean and median localization error of 30 pixels and 4 pixels respectively.

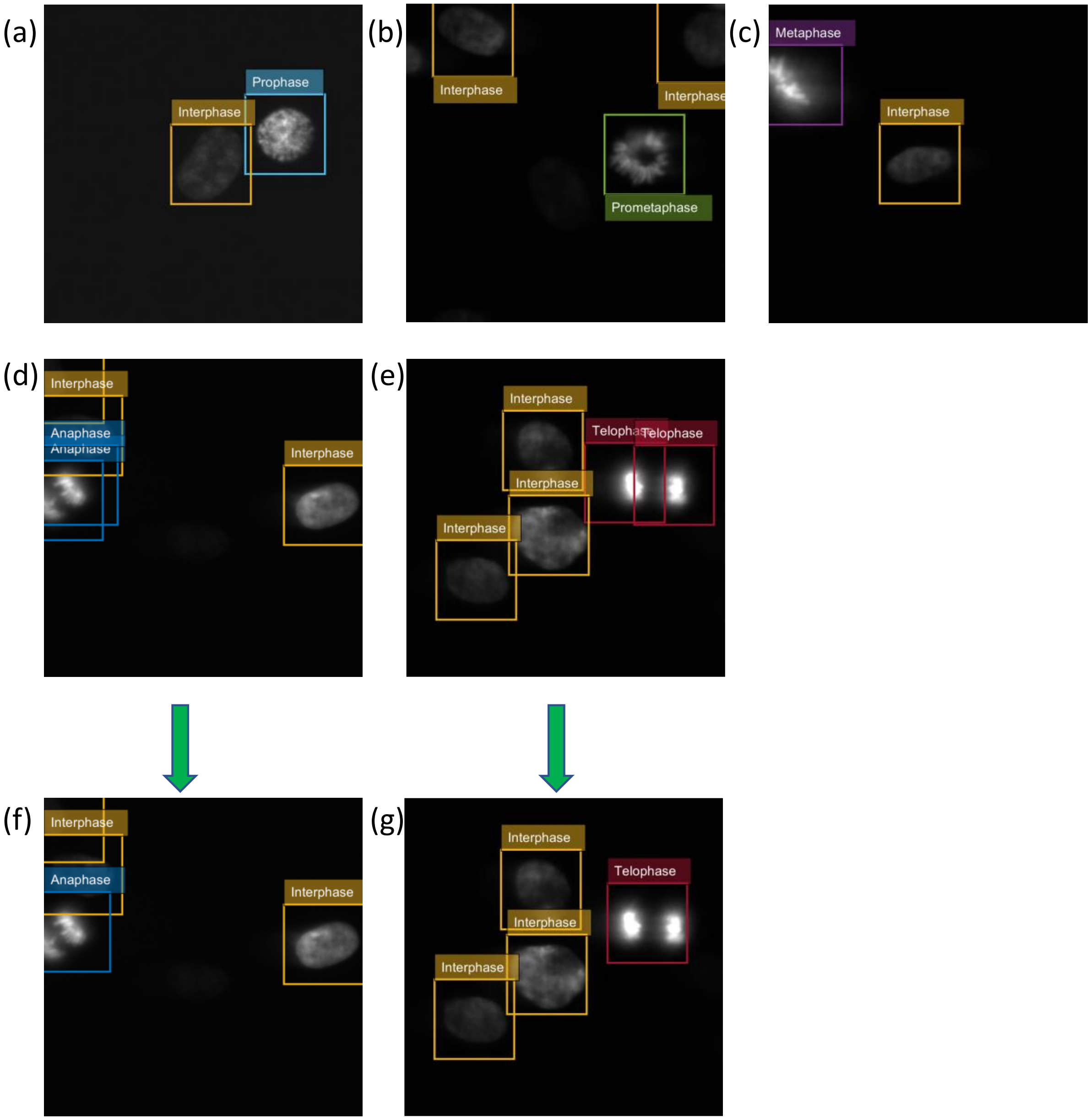


**Figure S10: Sample output from the YOLOv5 network.** Representative images showing (a) prophase, (b) prometaphase, (c) metaphase, (d) anaphase, and (e) telophase cells. Because the YOLO network occasionally will generate multiple detections for the two daughter cells in anaphase and telophase, we grouped detected mitotic cells within a radius of 10 µm as in (f) and (g).
